# Robust emergence of sharply tuned place cell responses in hippocampal neurons with structural and biophysical heterogeneities

**DOI:** 10.1101/681189

**Authors:** Reshma Basak, Rishikesh Narayanan

## Abstract

Hippocampal pyramidal neurons sustain propagation of fast electrical signals and are electrotonically non-compact structures exhibiting cell-to-cell variability in their complex dendritic arborization. In this study, we demonstrate that sharp place-field tuning and several somato-dendritic functional maps concomitantly emerge despite the presence of geometrical heterogeneities in these neurons. We establish this employing an unbiased stochastic search strategy involving thousands of models that spanned several morphologies and distinct profiles of dispersed synaptic localization and channel expression. Mechanistically, employing virtual knockout models, we explored the impact of bidirectional modulation in dendritic spike prevalence on place-field tuning sharpness. Consistent with prior literature, we found that across all morphologies, virtual knockout of either dendritic fast sodium channels or *N*-methyl-D-aspartate receptors led to a reduction in dendritic spike prevalence, whereas *A*-type potassium channel knockouts resulted in a nonspecific increase in dendritic spike prevalence. However, place-field tuning sharpness was critically impaired in all three sets of virtual knockout models, demonstrating that sharpness in feature tuning is maintained by an intricate balance between mechanisms that promote and those that prevent dendritic spike initiation. From the functional standpoint of the emergence of sharp feature tuning and intrinsic functional maps, within this framework, geometric variability was compensated by a combination of synaptic democracy, the ability of randomly dispersed synapses to yield sharp tuning through dendritic spike initiation, and ion-channel degeneracy. Our results suggest electrotonically non-compact neurons to be endowed with several degrees of freedom, encompassing channel expression, synaptic localization and morphological micro-structure, in achieving sharp feature encoding and excitability homeostasis.

## INTRODUCTION

Structure-function relationships are central to several biological research paradigms. From the standpoint of cellular neurophysiology, two major factors that impact neuronal function are its morphology and the ion channels and receptors that are expressed throughout neuronal membranes. Neuronal morphology has been shown to have strong links with its physiology, spanning a wide variety of functional characteristics, including neuronal excitability, firing patterns, intraneuronal coupling, intrinsic functional maps, somatodendritic subthreshold resonance, back- and forward propagation of electrical potentials and frequency selectivity in firing (Vetter et al. 2001; Mainen and Sejnowski 1996; Stiefel and Sejnowski 2007; Ostojic et al. 2015; Cannon et al. 2010; van Elburg and van Ooyen 2010; van Ooyen et al. 2002; Narayanan and Chattarji 2010; Narayanan et al. 2005; Dhupia et al. 2014; Krichmar et al. 2002; Ferrante et al. 2013; Beining et al. 2017; Weaver and Wearne 2008; Schaefer et al. 2003). Neuronal morphology has also been viewed to be a critical cog in solving the wiring optimization problem, which minimizes the cost of space, time, and matter (Cajal 1992; Cuntz et al. 2010; Cherniak 1992; Kim et al. 2012) towards transmitting input signals most efficiently (Chklovskii 2004). However, studies employing multicompartmental neuronal models have also demonstrated a remarkable degree of flexibility in a neuron’s ion channel distributions that can combine to elicit similar physiological outcomes (Rathour and Narayanan 2014; Basak and Narayanan 2018b; Otopalik et al. 2017b; Otopalik et al. 2017a; Otopalik et al. 2019; Migliore et al. 2018; Beining et al. 2017; Taylor et al. 2009).

In stomatogastric ganglion (STG) neurons, neuronal physiology has been shown to remain comparable irrespective of animal-to-animal variability in morphologies, introducing the concept of morphological sloppiness, whereby disparate morphologies endowed with significant heterogeneities yielded similar functional outcomes (Otopalik et al. 2017b; Otopalik et al. 2017a; Otopalik et al. 2019). The sloppy nature of morphological heterogeneities was shown to be reliant on these neurons being electrotonically compact, and on the slow temporal dynamics of the network that they reside in (Otopalik et al. 2017b; Otopalik et al. 2017a; Otopalik et al. 2019). However, electronic compactness or slow temporal dynamics of underlying signals are typically not characteristic features of mammalian excitatory neurons. For instance, hippocampal CA1 neurons have been shown to be electrotonically non-compact (Mainen et al. 1996; Carnevale et al. 1997), both due to passive cable filtering (Rall 1977; Stuart and Spruston 1998; Koch et al. 1983; Golding et al. 2005; Rall 1967) and substantial non-linearities in signal integration by active dendrites (Stuart and Sakmann 1994; Johnston et al. 1999; Hoffman et al. 1997; Spruston et al. 1995; Magee and Cook 2000; Narayanan and Johnston 2007, 2008, 2012; Losonczy and Magee 2006; Gasparini et al. 2004; Golding and Spruston 1998; Johnston and Narayanan 2008). These neurons sustain propagation of fast electrical signals, including back propagating action potentials (Spruston et al. 1995; Migliore et al. 1999; Golding et al. 2001; Johnston et al. 1999; Hoffman et al. 1997) and dendritic spikes (Golding and Spruston 1998; Gasparini et al. 2004; Losonczy and Magee 2006; Golding et al. 1999) and effectuate several forms of precise coincidence detection in sustaining baseline physiology and plasticity (Katz et al. 2007; Migliore 2003; Das and Narayanan 2015, 2017; Spruston 2008; Bittner et al. 2015; Bittner et al. 2017; Hoffman et al. 1997; Losonczy and Magee 2006; Magee and Johnston 1997; Takahashi and Magee 2009; Golding et al. 2002; Das et al. 2017). Is morphological heterogeneity in CA1 pyramidal neurons (Ambros-Ingerson and Holmes 2005) a strongly constraining stringent parameter or a sloppy parameter with reference to its impact on specific aspects of neuronal physiology? In STG neurons, electrotonic compactness compensates for geometric variability across animals, aided by the slow kinetics of the underlying signals. What strategy should electrotonic non-compact neurons endowed with active dendrites, with a need to sustain fast electrical signals across the somatodendritic arbor, follow to compensate for morphological heterogeneity across neurons?

Against this backdrop, we recently showed that the concomitant emergence of sharp place field tuning and intrinsic functional maps is resilient to heterogeneities in profiles of dispersed synaptic localization and ion channel expression in CA1 pyramidal neurons (Basak and Narayanan 2018b). We showed this to be consequent to the ability of randomly dispersed synapses to elicit dendritic spikes, resulting in precise transmission of temporal information along an active dendritic arbor. Can dendritic spikes occur with dispersed synaptic localization in different morphologies endowed with geometrical variability? How dependent is the concomitant emergence of sharp place-field tuning and intrinsic functional maps on morphological heterogeneities? Does the requirement for the propagation of fast events (dendritic spikes and backpropagating action potentials) make morphological heterogeneities to be a strongly constraining parameter in the emergence of sharp feature tuning? Can several intrinsic functional maps emerge together in the presence of morphological heterogeneities? What is the impact of bidirectional modulation of dendritic spike prevalence on sharpness of feature tuning?

To answer these questions, we studied place cell tuning in five distinct morphologically realistic CA1 neuron reconstructions, endowed with the same set of ion channel subtypes using a conductance-based framework. We used a stochastic search algorithm to generate thousands of unique models for each morphology, with identical parametric search space with identical pseudo-random sequences defining model parameters that spanned passive and active properties across morphologies. Neuronal models built from each morphological reconstruction received identical place-field inputs through 100 stochastically-activated excitatory synapses that were randomly dispersed across the *stratum radiatum* of the respective dendritic arbor. Together, the five distinct reconstructions accounted for the structural heterogeneities, whereas the random dispersion of place-field synapses and the disparate ion channel distribution profiles represented biophysical heterogeneities. This experimental strategy ensured that the parametric search space was unbiased and identical across all morphologies, that there was no specification of precise synaptic localization and allowed us to explore sharp tuning and intrinsic functional maps across a large search space involving structural and biophysical heterogeneities that are observed in CA1 pyramidal neurons.

We found that all morphologies yielded several unique models with disparate channel conductances and with randomly dispersed synaptic localization profiles, which had similar sharply tuned place cell response profiles and concomitantly expressed several intrinsic functional maps. The sharp tuning was effectuated by the generation of dendritic spikes, in response to place-field inputs arriving through randomly dispersed non-clustered synapses. We assessed the impact of bidirectional modulation of dendritic spike prevalence by virtual knockout of dendritic fast sodium (dNaF) channels or *N*-Methyl-D-Aspartate (NMDA) receptors (for reduction in the prevalence) or transient *A*-type potassium (KA) channels (for enhancing the prevalence). We found sharpness of place cell responses to substantially decrease in the absence of any of these components, either through a reduction in overall depolarizing drive (dNaF channels or NMDA receptors) or a non-specific enhancement of excitability (KA channels).

From the functional standpoint of the emergence of sharp feature tuning and intrinsic functional maps, our results provide clear lines of evidence towards neuronal morphology being a sloppy parameter even in an electrotonically non-compact neuron that is required to sustain propagation of fast electrical signals. In this case, dispersed synaptic localization, synaptic democracy and ion channel degeneracy together compensated for neuron-to-neuron variability in geometry. The ability of an electrotonically non-compact neuron to elicit sharp feature tuning and concomitantly maintain several intrinsic functional maps with disparate ion channel combinations in distinct morphologies and in the presence of randomly dispersed iso-feature synapses, provides a general strategy for neurons to ensure resilience to biophysical and morphological heterogeneities. The ability to elicit precise function with randomly dispersed synapses and in the face of morphological heterogeneities also precludes the need for *precise* wiring specificity of iso-feature synapses to specific locations on a neuronal morphology, especially in the hippocampus where place-cell formation manifests in an activity-dependent manner.

## METHODS

The computational methods and analysis protocols employed here were adopted from (Basak and Narayanan 2018b), and are listed below.

### Morphologies: Structure, passive and active properties

Five different morphologically realistic reconstructions of hippocampal CA1 neurons (Fig. 1a), namely, *n123*, *n128* (Pyapali et al. 1998), *n172*, *n175* and *n184* (Pyapali and Turner 1996) were obtained from the NeuroMorpho Database (Ascoli et al. 2007). Performing 3-dimensional Sholl’s analysis (Sholl 1953; Vyas et al. 2002; Narayanan et al. 2005) on these morphologies revealed heterogeneities in their overall morphology, the dendritic length and branching pattern (Fig. 1b). The base parameter values constituting the active and passive properties were the same for all the morphologies and were taken from (Basak and Narayanan 2018b; Rathour and Narayanan 2014). The specific membrane capacitance for all morphologies was set uniformly at 1 µF/cm^2^. *R_a_* and *R_m_* were set in a gradient along the trunk as a function of the radial distance of the compartment from the soma according to the equations and parametric values in Tables 1–2. The apical obliques had the same *R_a_* and *R_m_*as that of the trunk compartment from which they originated. The basal dendrites and the axonal compartments (only present in *n123*) had somatic *R*_m_ and *R*_a_. Each morphology was compartmentalized according to the *d*_λ_ rule (Carnevale and Hines 2006) to ensure isopotentiality in each compartment. Specifically, each compartment was smaller than 0.1×λ_100_, with λ_100_ representing the space constant of the section computed at 100 Hz.

**Figure 1.**
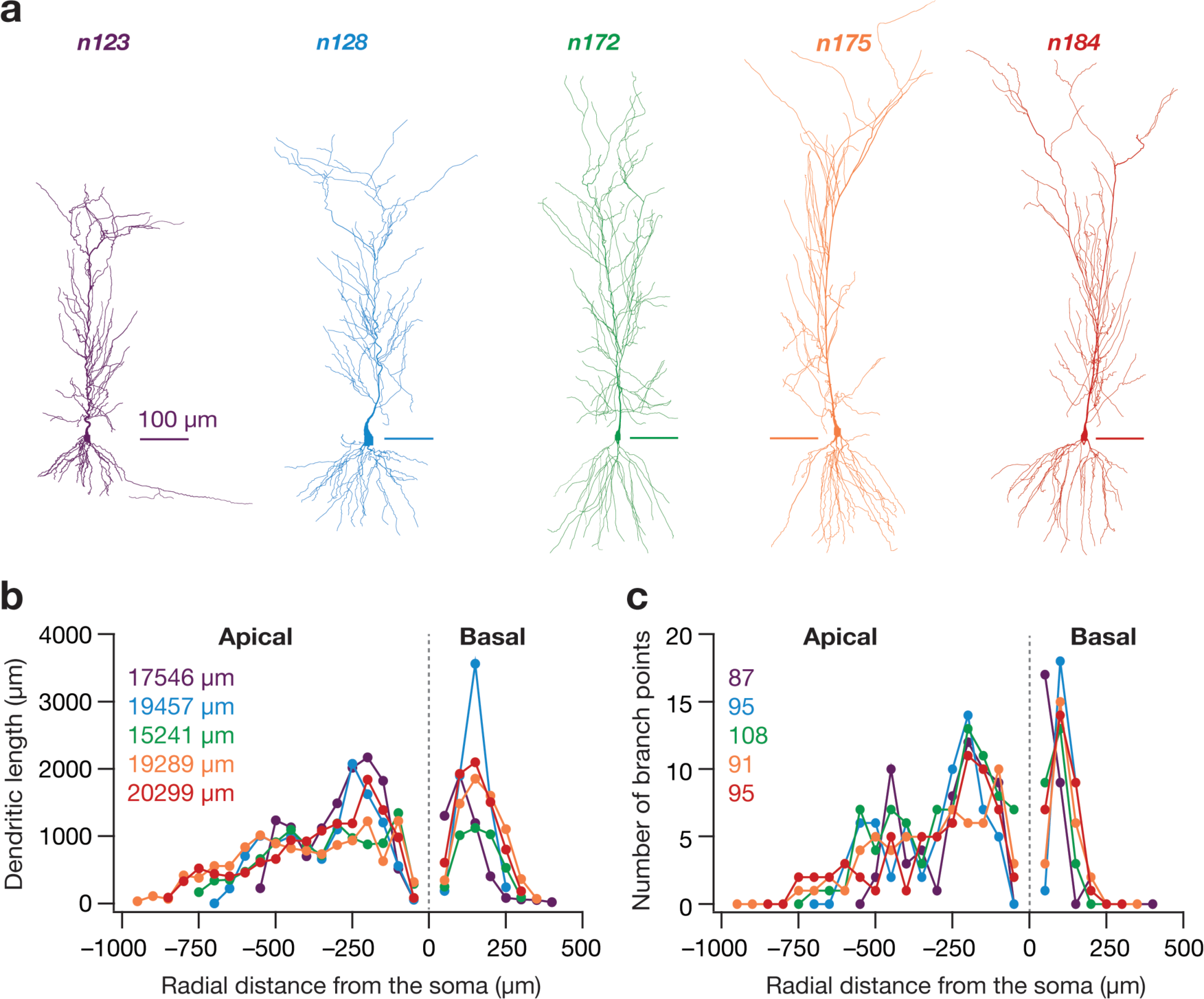
The five hippocampal pyramidal neuron morphologies and their properties. **a.** 2-D projections of the five different 3-D reconstructed hippocampal CA1 pyramidal neurons used in the study. Scale bars are 100 µm in length. **b.** Dendritic length along apical and basal dendritic axes of each morphology shown in (**a**), plotted as a function of radial distance from soma obtained through Sholl’s analysis. The numbers in inset correspond to the total dendritic length of each morphology. The color-codes corresponding to each morphology in (**a**) are consistent for all panels of this figure. **c.** The number of branching points in the apical and basal dendritic trees of each morphology shown in (**a**) as a function of radial distance from the soma, obtained through Sholl’s analysis.

**Table 1.** Description of somatodendritic gradients in passive properties and channel gradients used in the model. *x* represents radial distance from the soma. These distributions are identical to those described in (Basak and Narayanan 2018b).

| Parameter | Gradient of distribution |
| --- | --- |
| Specific membrane resistivity, $R_m$ | $R_m(x) = R_m + \frac{(R_m - \text{min} - R_m - \text{max})}{1 + \exp((R_m - \text{hmp} - x) / R_m - \text{slope})} \text{k}\Omega / \text{cm}^2$ |
| Axial resistivity, $R_a$ | $R_a(x) = R_a + \frac{(R_a - \text{min} - R_a - \text{max})}{1 + \exp((R_a - \text{hmp} - x) / R_a - \text{slope})} \Omega.\text{cm}$ |
| Maximal conductance of NaF channels | $\bar{g}_{\text{Na}}$ , uniform across the somatodendritic arbor, with the density at the axon initial segment set at $5 \bar{g}_{\text{Na}}$ |
| Maximal conductance of KDR channels | $\bar{g}_{\text{KDR}}$ , uniform across the somatodendritic arbor |
| Maximal conductance of KA channels | $g_{\text{KA}}(x) = \bar{g}_{\text{KA}} \left( 1 + \frac{\bar{g}_{\text{KA}} - \text{fold}}{100} \right) \text{mS}/\text{cm}^2$ |
| Maximal conductance of $h$ channels | $g_h(x) = \bar{g}_h \left( 1 + \frac{\bar{g}_h - \text{fold}}{1 + \exp((\bar{g}_h - \text{hmp} - x) / \bar{g}_h - \text{slope})} \right) \mu\text{S}/\text{cm}^2$ |
| Maximal conductance of CaT channels | $g_{\text{CaT}}(x) = \bar{g}_{\text{CaT}} \left( 1 + \frac{\bar{g}_{\text{CaT}} - \text{fold}}{1 + \exp((\bar{g}_{\text{CaT}} - \text{hmp} - x) / \bar{g}_{\text{CaT}} - \text{slope})} \right) \mu\text{S}/\text{cm}^2$ |

**Table 2.**
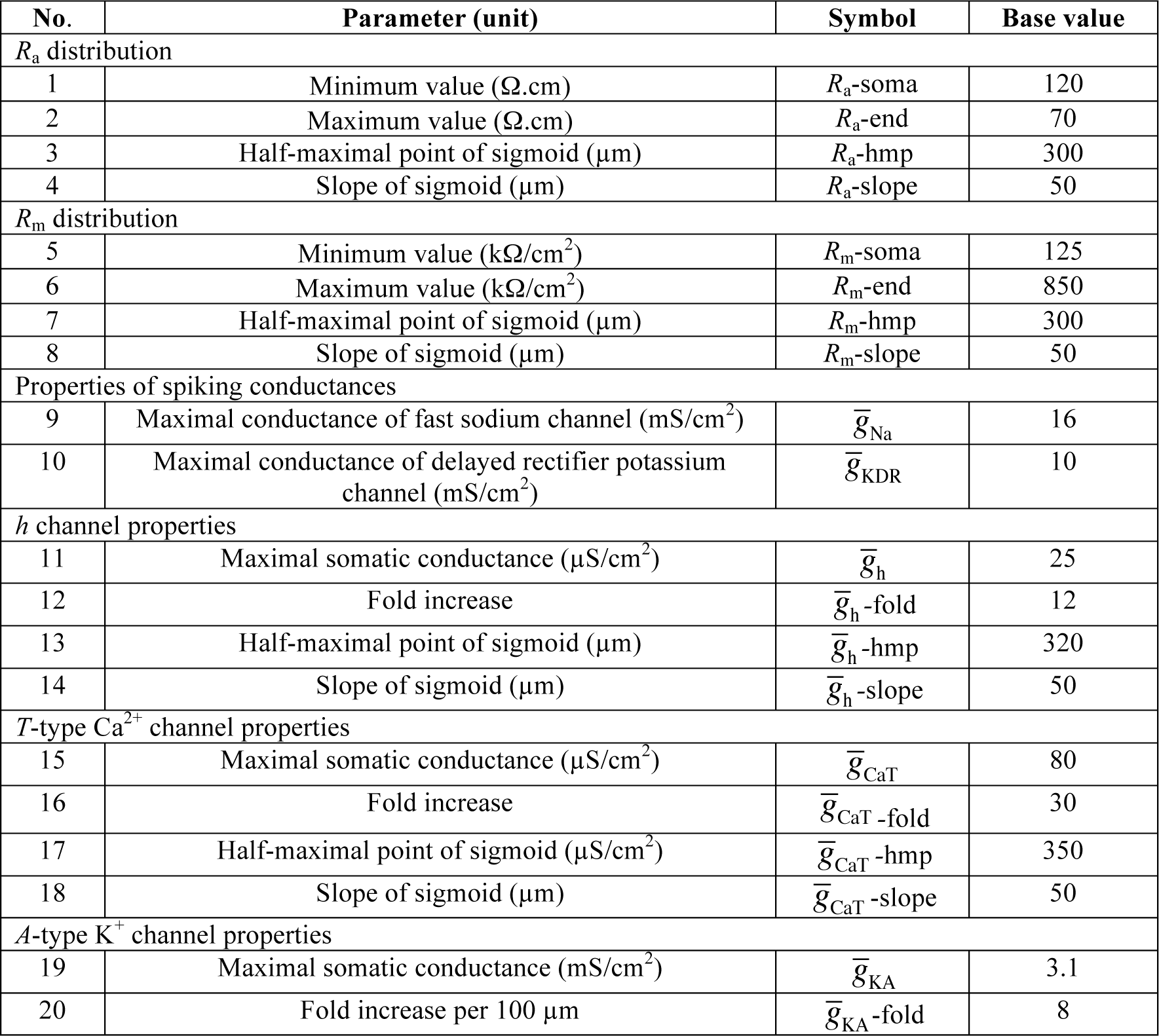
Model parameters and their base values for the model. These parameter values are identical to those described in (Basak and Narayanan 2018b).

Five different ion channels were incorporated into all morphologies (Basak and Narayanan 2018b). These were: Hodgkin-Huxley-type delayed rectifier potassium (KDR), fast sodium (NaF), *T-*type calcium (CaT), hyperpolarization-activated cyclic-nucleotide-gated (HCN) nonspecific cation and *A*-type potassium (KA) channels. Currents through the NaF, KDR, KA and HCN channels used an Ohmic formulation with reversal potentials for Na^+^, K^+^ and *h* channels set at 55, –90 and –30 mV respectively. The CaT current was modeled using the Goldman-Hodgkin-Katz (GHK) formulation (Shah et al. 2008). NaF and KDR conductances were distributed uniformly in the soma and across the dendritic arbor with respective maximal conductances set at 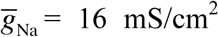 and 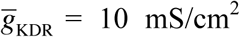 (Magee and Johnston 1995; Hoffman et al. 1997). As the recovery of dendritic sodium channels from inactivation is slower (Colbert et al. 1997), an additional inactivation gating variable was included in the model for Na^+^ channels that expressed in the apical dendrites (Migliore et al. 1999; Magee and Johnston 1995). The three subthreshold channel conductances (CaT, HCN and KA) were distributed with increasing somato-apical gradients (Table 1–2). Two different KA conductance models were used for proximal (≤100 µm radial distance from soma) and distal (>100 µm) apical dendritic compartments (Migliore et al. 1999). The half-maximal activation voltage for HCN channels was –82 mV for proximal apical compartments (radial distance ≤ 100 µm), linearly varied from –82 mV to –90 mV for compartments between 100 and 300 µm, and was set at –90 mV for compartments with distances larger than 300 µm (Magee 1998). All active and passive properties of basal dendritic compartments for all morphologies were set to their respective somatic values.

It is critical to note here that the intrinsic parametric values and distribution in each morphology was *not* tuned to yield a physiologically plausible base model with functional maps (Narayanan and Johnston 2012) matching experimental values. Rather, we chose to employ a stochastic search approach to find models with disparate parametric combinations, which yielded physiologically relevant intrinsic CA1 pyramidal cell properties (refer to the section “Multiparametric multi-objective stochastic search algorithm” below). This approach ensured that our conclusions were not biased by the narrow parametric regimes of a single hand-tuned model, but account for biological heterogeneities in the biophysical properties of hippocampal neurons (Foster et al. 1993; Goldman et al. 2001; Prinz et al. 2004; Marder and Taylor 2011; Rathour and Narayanan 2012, 2014; Basak and Narayanan 2018b).

### Synapses: Models, localization and permeabilities

Colocalized N-Methyl-D-Aspartate (NMDA) and α-amino-3-hydroxy-5-methyl-4-isoxazolepropionic acid (AMPA) receptors constituted a single synapse (Basak and Narayanan 2018b). Both receptors were modeled using the GHK formulation with the NMDAR:AMPAR ratio set at 1.5. The kinetics of AMPA and NMDA receptor currents were adopted from previous literature (Narayanan and Johnston 2010; Ashhad and Narayanan 2013; Anirudhan and Narayanan 2015). The current through the NMDA receptor, as a function of voltage and time, was dependent on three ions: sodium, potassium and calcium. Consequently, as per the Goldman-Hodgkin-Katz formulation:

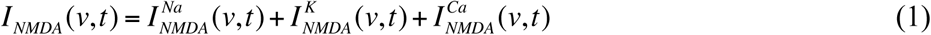

where,

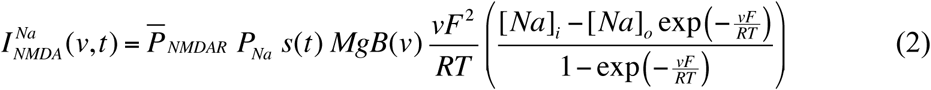

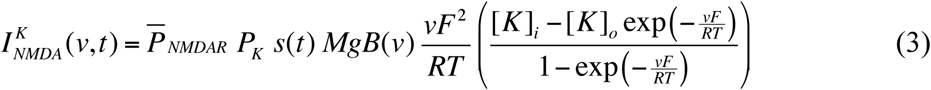

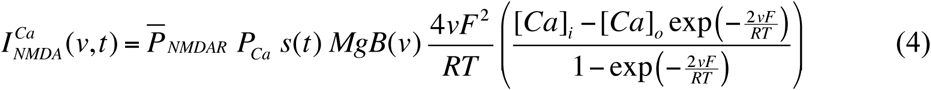

where 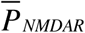 is the maximum permeability of the NMDA receptor. The relative permeability ratios were set at *P_Ca_*=10.6, *P_Na_* =1, *P_K_* =1. Default values of concentrations were (in mM): [*Na*]_i_=18, [*Na*]_o_=140, [*K*]_i_=140, [*K*]_o_=5, [*Ca*]_i_=100 × 10^−6^, [*Ca*]_o_=2. *MgB(v)* governed the magnesium dependence of the NMDAR current, given as (Jahr and Stevens 1990):

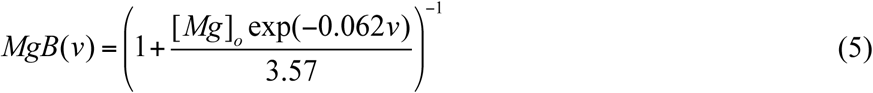

with the default value of [*Mg*]*_o_*set at 2 mM. *s*(*t*) governed the kinetics of the NMDAR current, and is given as:

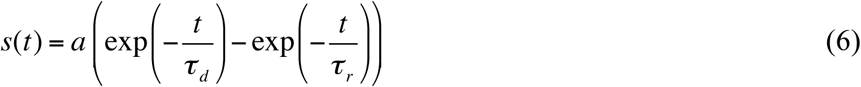

where *a* is a normalization constant, making sure that 0 ≤ *s*(*t*) ≤ 1, *τ_d_* is the decay time constant, *τ_r_* is rise time, with *τ_r_* =5 ms, and default *τ_d_* =50 ms (Narayanan and Johnston 2010; Ashhad and Narayanan 2013).

Current through the AMPA receptor was modeled as the sum of currents carried by sodium and potassium ions:

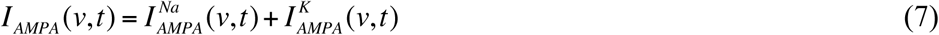

where,

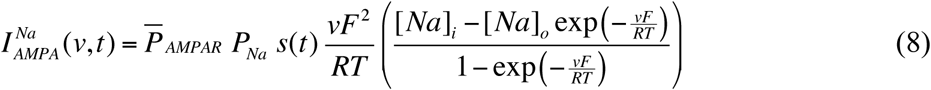

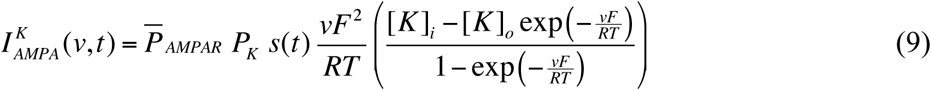

where 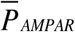 is the maximum permeability of the AMPA receptor. The relative permeability ratios *P_Na_* and *P_K_* were equal and set to 1. *s*(*t*) was the same as that for the NMDA receptor, but with τ_r_=2 ms and τ_d_=10 ms (Narayanan and Johnston 2010). AMPAR permeabilities for synapses for all somato-apical locations, for all morphologies (Fig. 2c) were adjusted such that the unitary somatic response amplitude was ∼0.2 mV (Fig. 2d) irrespective of synaptic location (Andrasfalvy and Magee 2001; Magee and Cook 2000). This *synaptic democracy* (Hausser 2001) ensured that attenuation along the dendritic cable did not play a critical role in determining the impact of synaptic localization profiles on tuning properties, also precluding the requirement of explicitly modeling spines. We have previously established that dispersed synaptic localization yields sharp place field tuning(Basak and Narayanan 2018b). Consequently, we randomly dispersed 100 synapses on the somato-apical dendritic arbor within the *stratum radiatum* of each morphology, also ensuring that there was no spurious clustering as a consequence of such random dispersion (Fig. 2a–b).

**Figure 2.**
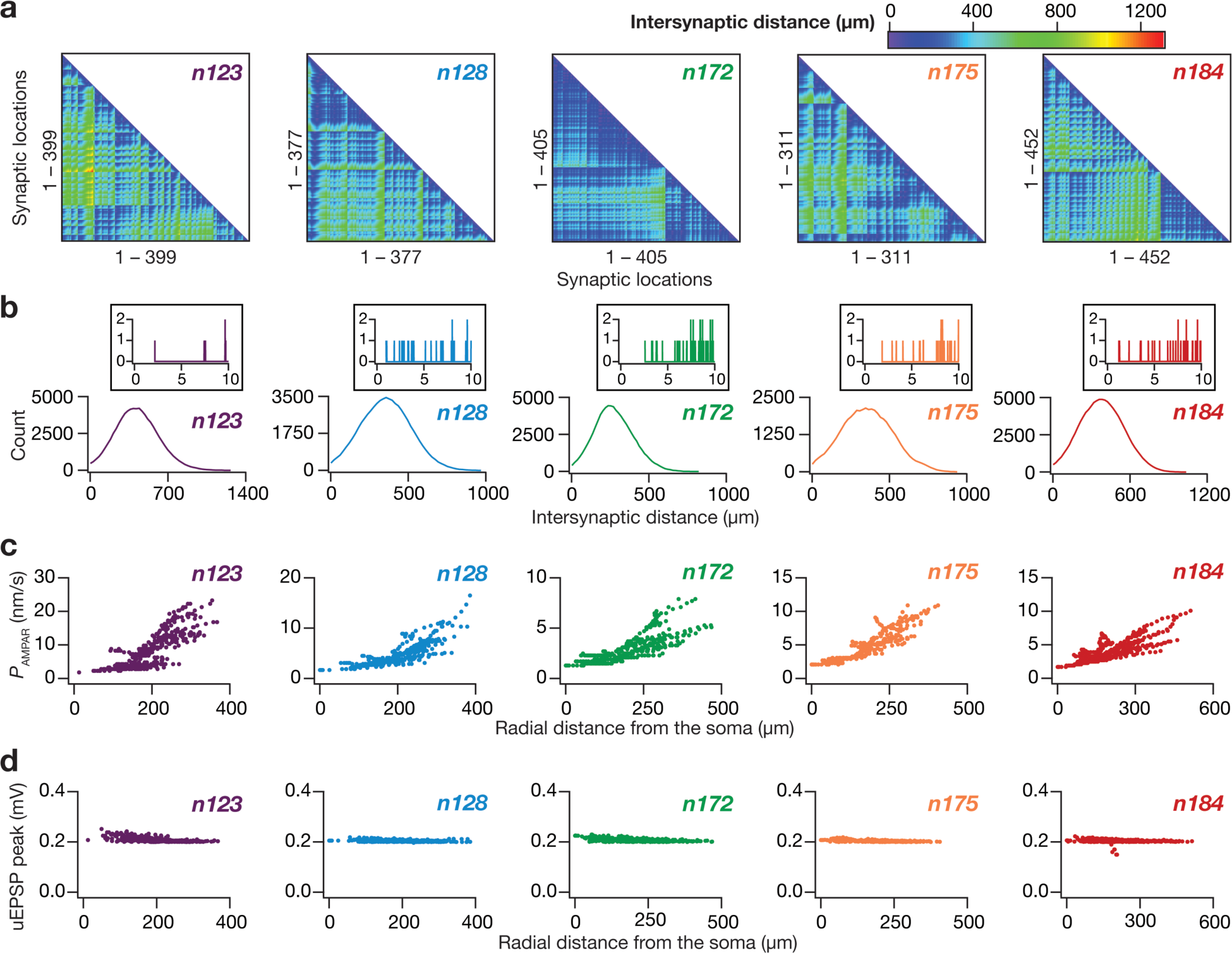
Synaptic localization and strength of randomly dispersed synapses in the five hippocampal pyramidal neuron morphologies. **a.** From left to right: lower triangular part of the intersynaptic distance matrix in models where synapses were randomly dispersed across the *stratum radiatum* of their respective apical dendritic arbor. Note that the first panel (morphology *n123*) is redrawn from Fig. 4B of (Basak and Narayanan, 2018), as the models for morphology *n123* were derived with synaptic locations as in (Basak and Narayanan, 2018). **b.** Distribution of distances plotted in the matrices given in panel (**a**). Inset, distribution of distances zoomed to 0– 10 µm, showing only a small proportion of synaptic pairs with intersynaptic distances less than 10 µm. **c.** From left to right: location dependent permeability values AMPA receptor that normalized somatic unitary postsynaptic potential (uEPSP) amplitudes to around 0.2 mV. **d.** From left to right: the resulting somatic uEPSP values from AMPA receptor permeabilities shown in (**c**). Note that all somatic uEPSP values cluster around 0.2 mV, irrespective of the somato-apical location of the synapse. For all panels, the synaptic properties are depicted for morphologies *n123*, *n128*, *n172*, *n175* and *n184* in that order.

### Place cell inputs and instantaneous firing rate analysis

A probability density function modeled as a Gaussian-modulated cosinusoid governed the excitatory drive of the impinging synapses on the place cell. The cosinusoid, whose frequency was set at 8 Hz, mimicked the theta activity present in hippocampus during exploratory and REM sleep stages, whereas the Gaussian function mimicked the increase in excitatory drive in the place cell as a result of the animal entering the place field (Geisler et al. 2010; Buzsaki 2002, 2006; Basak and Narayanan 2018b). Therefore, each synapse in the neuron received presynptic action potentials whose probability of occurrence at any given time point was defined by:

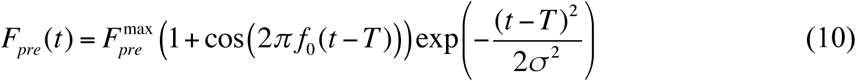

where *T* (5 s) defined the travel time between place field centers, *f*_0_ represented the cosine wave frequency (8 Hz), 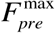 regulated the maximal input firing rate and σ defined the width of the Gaussian and controls the extent of the place field (1 s).

One hundred independent synapses receiving afferent inputs that obeyed this distribution drove postsynaptic place cell response, which were recorded as somatodendritic voltages which yielded somatic spike timings. These spike times were then digitized into binary values and then converted into instantaneous firing rate profiles by convolution with a Gaussian kernel. This smoothed instantaneous firing rate was representative of the place cell response (*e.g.*, Fig. 3a). Following this, two measurements were derived from the firing rate profiles to determine tuning sharpness of the response: (i) the maximum firing rate of place cell (*F*_max_); (ii) the full-width at half maximum (FWHM) of the profile. Cells having a firing profile with high *F*_max_ and low FWHM were considered to be sharply tuned to the place cell input. As the input distribution was fixed, this constituted a relative measure of tuning sharpness, comparing output responses of different neuronal models when similar inputs were presented. The place cells with firing rate profiles having FWHM ≤ 2.8 s and *F*_max_ ≥ 40 Hz were considered to have sharp place field tuning.

**Figure 3.**
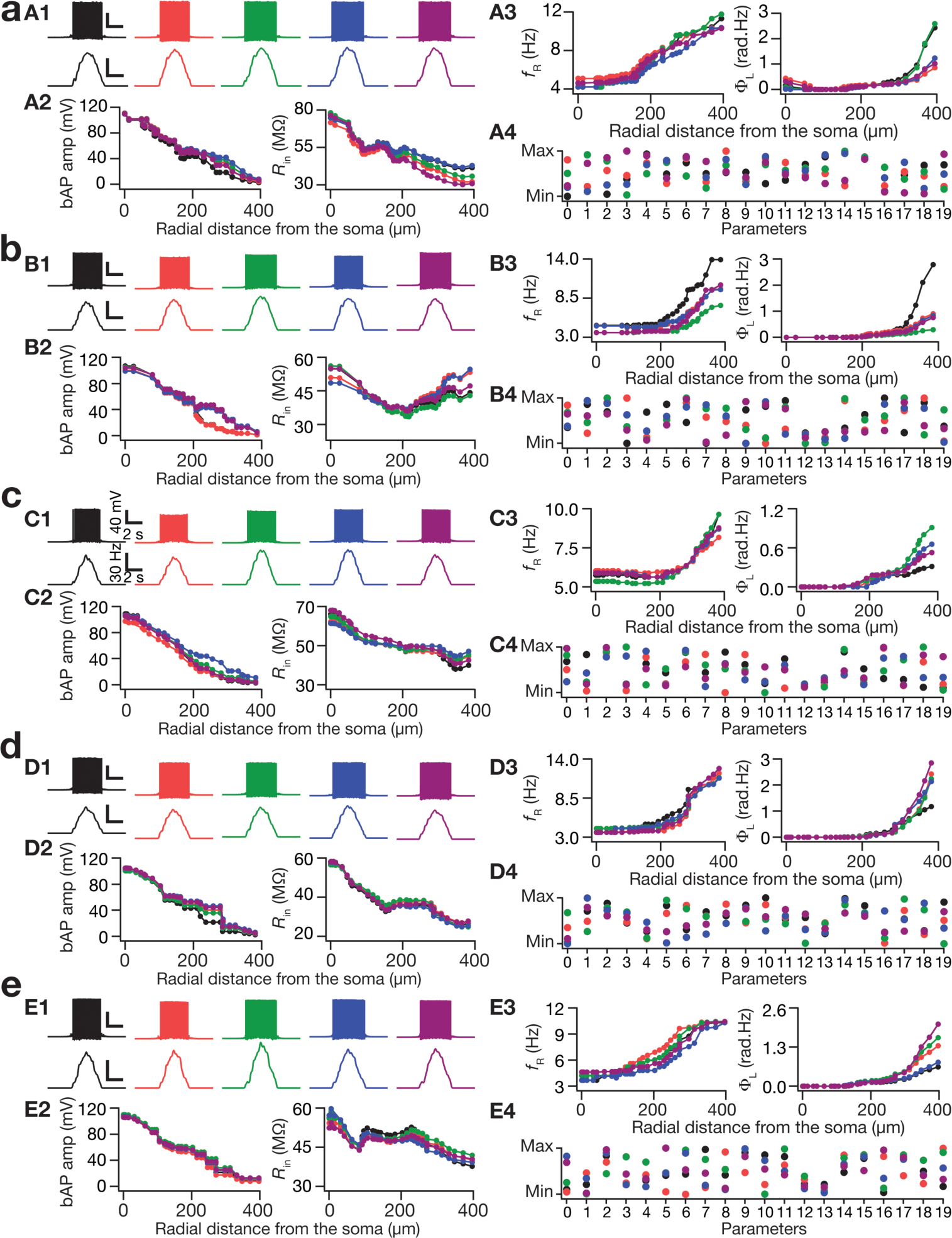
Expression of ion channel degeneracy in the concomitant emergence of sharp place-cell tuning and four functional maps in each of the five morphologies. **a. A1.** Five example valid model voltage traces (top), the corresponding firing rates (bottom) showing similar sharp tuning. Note that the normalized parameters span the entire available range, still yielding similar place field tuning. **A2, A3.** Functional maps along the somato-apical trunk of back propagating action potential (bAP) amplitude (**A2**, left), input resistance (*R*_in_; **A2**, right), resonance frequency (*f*_R_; **A3**, left) and total inductive phase (Φ_L_, **A3**, right) of the five valid models whose voltage traces are shown in (**A1**). **A4.** Normalized values of parameters underlying the five models whose physiological properties are depicted in panels (**A1–A4**). **b, c, d, e.** The results depicted in (**a**) were derived from simulations employing morphology *n123*. The results in **b**, **c**, **d**, **e** are depicted employing the same organization in (**a**), for morphologies *n128*, *n172*, *n175* and *n184*, respectively. Note that for morphology *n123*, the example five models were derived from the same valid model population presented in (Basak and Narayanan 2018b).

**Figure 4.**
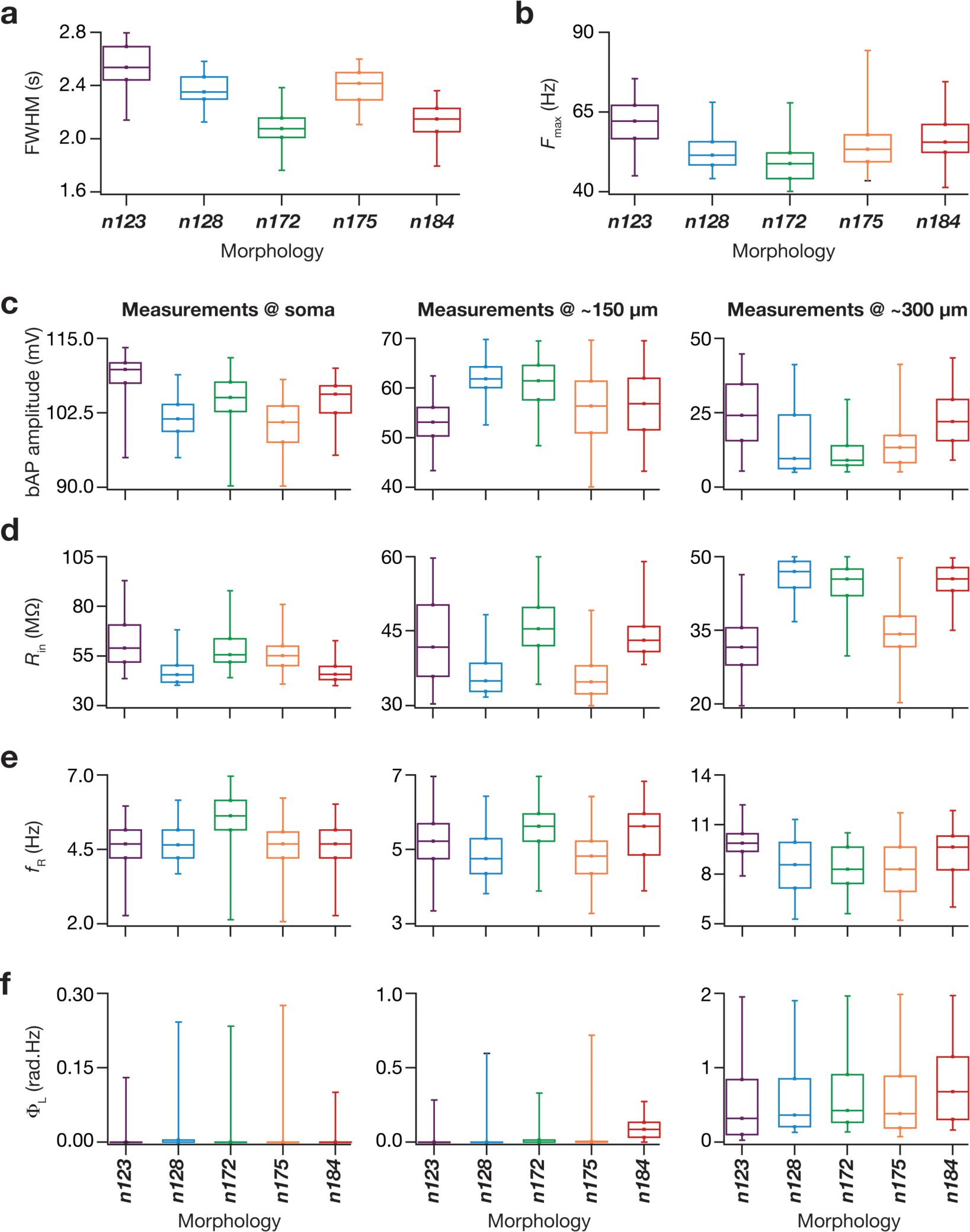
The valid model population dervied from simulations employing all five morphologies manifested sharpness of place-field tuning and matched intrinsic electrophysiological measurements from hippocampal pyramidal neurons. **a.** Box plots depicting the full-width at half-maximum (FWHM) values from the firing rate profiles of all the valid models from all five morphologies. The color-codes for this figure are consistent with that of the morphology colors depicted in Fig. 1a. **b.** Box plots depicting the maximum firing rate (*F*_max_) values of all the valid models from all five morphologies. **c.** Box plots depicting the bAP amplitude values of all the valid models from all five morphologies at soma (left), at 150 µm from soma (middle) and at 300 µm from soma (right). **d.** Box plots depicting the *R*_in_ values of all the valid models from all five morphologies at soma (left), at 150 µm from soma (middle) and at 300 µm from soma (right). **e.** Box plots depicting the *f_R_* values of all the valid models from all five morphologies at soma (left), at 150 µm from soma (middle) and at 300 µm from soma (right). **f.** Box plots depicting the Φ_L_ values of all the valid models from all five morphologies at soma (left), at 150 µm from soma (middle) and at 300 µm from soma (right). All box plots show medians and depict the four quartiles.

**Figure 5.**
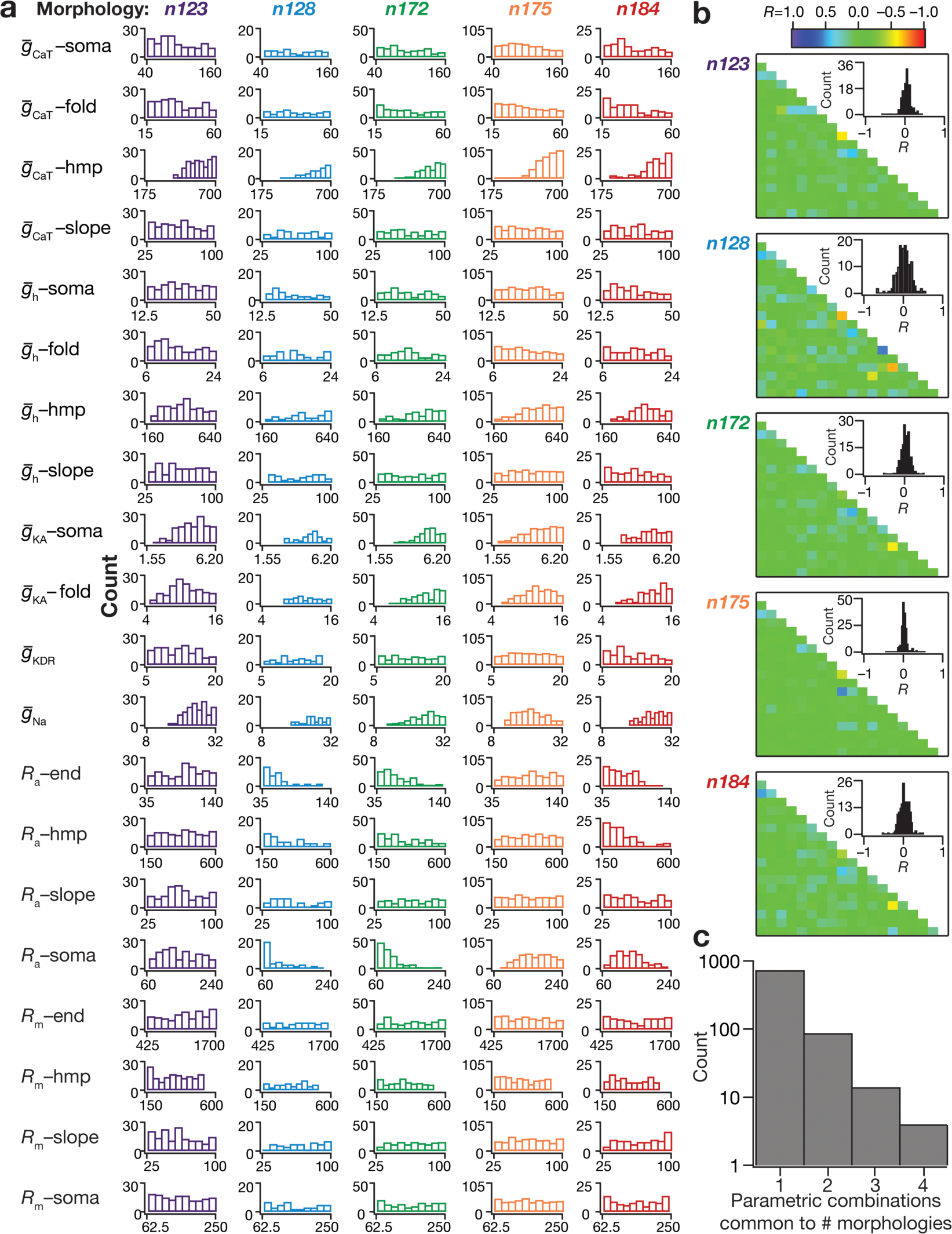
Across all morphologies, sharp place-cell tuning and intrinsic functional maps concomitantly emerged in the absence of strong pairwise correlations among underlying parameters. **a.** Histograms depicting the distribution of 20 underlying parameters constituting all the valid models from morphologies *n123*, *n128*, *n172*, *n175* and *n184* (left to right). **b.** From top to bottom: the pairwise parametric correlation coefficient (*R*) matrices of valid models from all five morphologies. Insets: Histograms, derived from the corresponding correlation coefficient matrices, depicting the pairwise parametric correlations of the valid models. **c.** Histogram depicting the number of parametric combinations that were common to valid models from different morphologies. The count corresponding to “1” depicts the number of parametric combinations that provided valid models only with a single morphology. The counts corresponding to “2–4” depict the number of parametric combinations that provided valid models with 2–4 morphologies, respectively. There was no parametric combinations that yielded valid models for all 5 morphologies.

### Multi-Parametric Multi-Objective Stochastic Search and Model Validation

In order to adequately explore comprehensive ranges of the critical parameters underlying our neuronal models and to test the robustness of our findings to parametric variability, we employed a Multi-Parametric Multi-Objective Stochastic Search (MPMOSS) approach (Foster et al. 1993; Goldman et al. 2001; Prinz et al. 2004; Marder and Taylor 2011; Rathour and Narayanan 2012, 2014; Anirudhan and Narayanan 2015; Mukunda and Narayanan 2017; Mishra and Narayanan 2019; Mittal and Narayanan 2018; Basak and Narayanan 2018b; Srikanth and Narayanan 2015).

To do this, we assigned a range to each of the 20 model parameters, which spanned 0.5×–2× of its physiological base value (Table 2) (Rathour and Narayanan 2014; Basak and Narayanan 2018b). For each trial, each parameter was independently and randomly chosen through uniform sampling of its respective range, which collectively constituted one model. 8000 (*n123*) – 10000 (*n128, n172, n175, n184*) independent models were generated for each morphology to explore the active and passive parametric space governed by the 20 model parameters (Table 2). Identical place cell inputs were fed to each of them through excitatory synapses and their individual somatic voltage responses and spike times were recorded to construct instantaneous firing rate profiles (see above). Specifically, although each of the 100 synapses impinging on a given morphology were independently sampled from the distribution in Eq. 10, individual synapses (*e.g.*, synapse 1 of each morphology had the same input pattern) of each morphology across models received the same input. This intra-model independence and inter-model equivalence was preserved for every synapse across models and across morphologies. These firing rate profiles were then subjected to two stages of validation. The first stage validated the model on its *encoding characteristics*, with high tuning sharpness (low FWHM and high *F*_max_) constituting the validation criteria. The second stage of validation assessed whether models from each morphology, already validated for sharp tuning of place cell responses, also matched *signature electrophysiological characteristics* of hippocampal neuronal somata and dendrites, as described below.

### Intrinsic measurements and model validation

To assess which of the sharply-tuned models exhibited signature physiological characteristics of hippocampal CA1 neurons, we subjected these models to a further step of validation involving four intrinsic functional maps (*e.g.*, Fig. 3A2–A3), namely: back-propagating action potentials (bAP), input resistance (*R*_in_), resonance frequency (*f*_R_) and total inductive phase (Φ_L_). These measurements were computed for three somatoapical trunk locations (at soma, at ∼150 µm radial distance from soma, at ∼300 µm radial distance from soma) for all the sharply tuned models in all five morphologies using protocols described in previous literature (Narayanan and Johnston 2007; Rathour and Narayanan 2014; Hoffman et al. 1997; Narayanan and Johnston 2008; Spruston et al. 1995). Briefly, to measure dendritic bAP, an action potential was initiated at the soma (2 nA current for 1 ms) and the bAP amplitude was measured at the three chosen locations along the somatoapical trunk. *R*_in_ was measured by injecting subthreshold current pulses of amplitudes spanning –50 pA to +50 pA, in steps of 10 pA and recording the local voltage responses to these current pulses. The respective steady state voltage responses at a given location were plotted against the corresponding current amplitudes to obtain the *V*-*I* plot. The slope of a linear fit to this steady-state *V*-*I* plot was taken as the *R*_in_ for that location, and the procedure was repeated at the three chosen locations along the somatoapical trunk.

Impedance-based measurements of the model were computed by injecting a chirp stimulus (Narayanan and Johnston 2007, 2008; Rathour and Narayanan 2012, 2014): a sinusoidal current wave with constant amplitude (100 pA; peak-to-peak) with frequency linearly increasing from 0.1 to 15 Hz in 15 s. The Fourier transform of the local voltage response was divided by the Fourier transform of the chirp stimulus to obtain the complex valued impedance *Z*(*f*), as a function of frequency *f*. The impedance amplitude profile |*Z*(*f*)| was computed as the magnitude of this impedance:

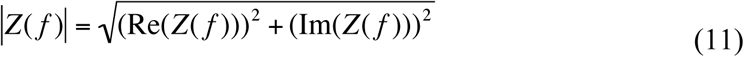

where Re(*Z*(*f*)) and Im(*Z*(*f*)) were the real and imaginary parts of the impedance |*Z*(*f*)|, respectively. The frequency at which |*Z*(*f*)| reached its maximum value was measured as the resonance frequency, *f*_R_, and was computed at the three chosen locations along the somatoapical trunk. The impedance phase profile *ϕ*(*f*) was computed as:

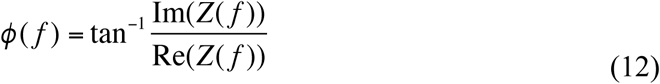

Φ_L_, defined as the area under the inductive part of *ϕ*(*f*), (Narayanan and Johnston 2008), was computed at the chosen locations along the somato-apical trunk based on the local impedance phase profile:

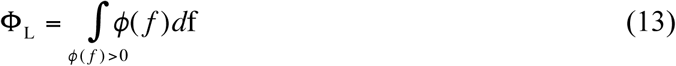

A total of 12 (4 intrinsic measurements × 3 somatoapical locations) constraints were imposed on the models showing sharp tuning across all morphologies. Neuronal models whose measurements fell within the values specified in Table 3 were considered to be valid. Note that for morphology *n123*, the valid model population from (Basak and Narayanan 2018b) has been reused; the other morphological reconstructions were not employed in (Basak and Narayanan 2018b), and were assessed as part of this study.

**Table 3.** Bounds on 12 measurements for the model to be intrinsically valid. These measurement bounds are identical to those described in (Basak and Narayanan 2018b), and are derived from electrophysiological measurements from CA1 pyramidal neurons (Spruston et al. 1995; Narayanan and Johnston 2007, 2008, 2010; Malik et al. 2016).

| Measurement | Soma | ~150 $\mu\text{m}$ | ~300 $\mu\text{m}$ |
| --- | --- | --- | --- |
| bAP Amplitude (mV) | 90–115 | 40–70 | 5–45 |
| Input resistance, $R_{\text{in}}$ ($\text{M}\Omega$) | 40–100 | 30–60 | 10–50 |
| Resonance frequency, $f_{\text{R}}$ (Hz) | 2–7 | 3–7 | 5–14 |
| Total inductive phase, $\Phi_{\text{L}}$ (rad Hz) | 0–0.3 | 0–1 | 0.025–2 |

### Dendritic spikes and backpropagating action potentials

To analyze the dynamics of dendritic spikes (dSpikes) or back propagating action potentials (bAPs) in the apical dendritic tree from each valid model across each morphology, we recorded voltage traces from 5 different somatoapical trunk (Fig. 6a) locations (0 µm, 100 µm, 150 µm, 200 µm, 250 µm and 300 µm radial distance from soma) in these models. We computed the peak of the spike at these locations for each somatic action potential, detected at the upstroke of the somatic voltage crossing –20 mV. We then calculated the differences between the timings of the peak at different dendritic voltages and the timing of the peak somatic voltage according to the equation:

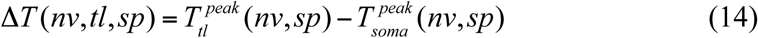

**Figure 6.**
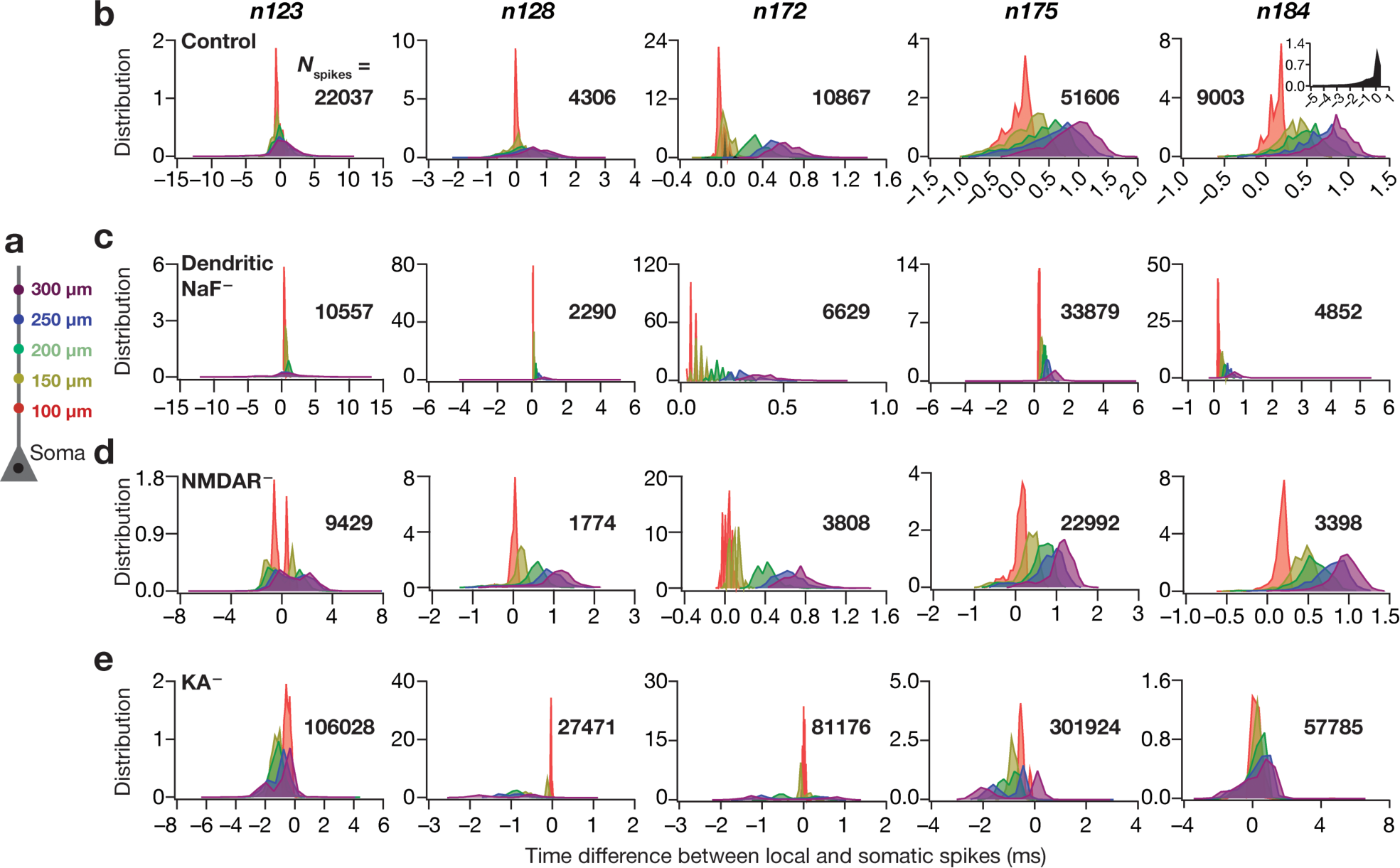
Virtual knockout of *A*-type potassium channels increased dendritic spike prevalence whereas knockouts of dendritic sodium channels or NMDA receptors decreased the prevalence. **a.** Schematic of the neuronal somato-apical trunk showing various points where the voltages were recorded. **b.** Left to right: histograms quantifying the differences between the timing of somatic action potential (AP) and bAP/dendritic spike (dSpike) along the somato-apical axis for all valid cells from each of the morphologies, namely *n123*, *n128*, *n172*, *n175* and *n184*. Note that negative temporal differences correspond to dSpikes which precede somatic AP, whereas positive values indicate bAPs which follow the somatic APs. The color-codes are derived from the schematic in (**a**), and depict the recording locations along the somato-apical trunk. Inset on far right panel: dendritic spikes were highly prevalent in an apical oblique of morphology *n184*. **c**–**e**. Same as (**b**), but for apical dendritic sodium channel (**c**; Dendritic NaF^−^), NMDA receptor (**d**; NMDAR^−^) or *A*-type potassium channel (**e**; KA^−^) knockouts. *N*_spikes_ refers to the number of spikes from different valid models belonging to the same morphology in plotting the corresponding distribution. Note that the data is represented as histograms, where the area under the curve for each plot is unity.

where Δ*T* represented the time difference, *nv* indexed the number of valid models (1, …, *N*_valid_), *tl* represented the five different trunk locations and *sp* represented the current somatic spike index, and spanned all the somatic spikes obtained for valid model number *nv*. 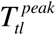 and 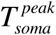 represented the time at which the spike reached its peak at location *tl* and at the soma respectively. This difference was computed for all spikes in all valid models, and was binned into the appropriate location *tl*. This was repeated for all the five morphologies to compare their dendritic spike dynamics.

A dendritic voltage peak preceding the somatic voltage peak (negative Δ*T*) was considered be dSpike at the recorded trunk location, whereas a dendritic peak following the somatic peak (positive ΔT) was representative of a bAP at that location. In order to quantify the fraction of dSpikes for each morphology, at each location we plotted normalized histograms of Δ*T* (Fig. 6b) and calculated the area under the curve (AUC) for the negative portion of Δ*T* axis (*e.g.*, Fig. 7a), for each location *tl*. This analysis was repeated for all virtual knockout models of all morphologies in order to quantify the effect of various channel conductances on dSpike dynamics.

**Figure 7.**
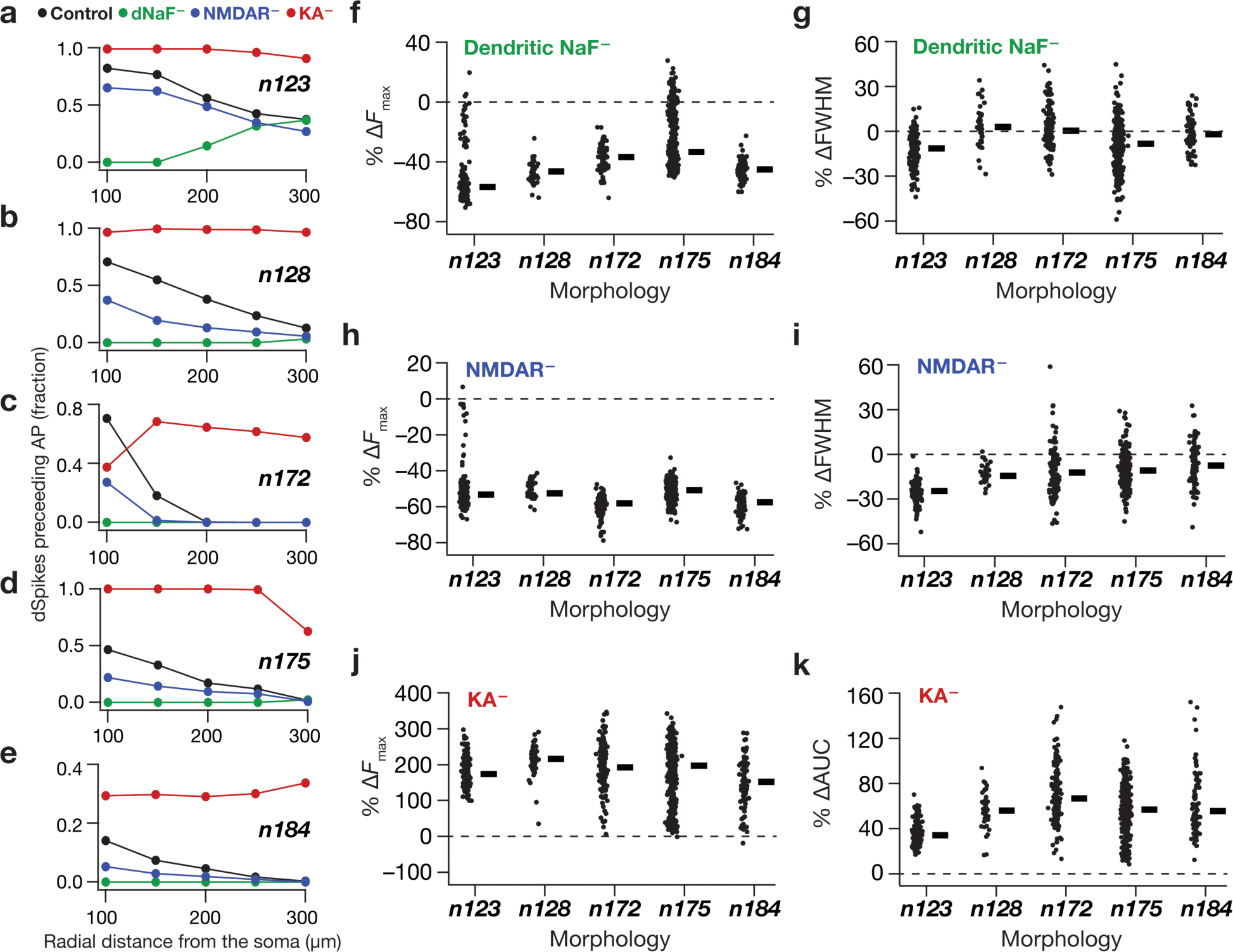
Tuning sharpness decreased in absence of dendritic sodium channels, NMDA receptors or transient potassium channels across all morphologies through bidirectional changes in dendritic spike generation. **a.** Plots, derived from histograms in (Fig 6. **b–e**), depicting the fraction of somatic APs where the dendritically recorded spike preceded the somatic AP for valid cells in *n123*. **b–e.** Same as (**a**) but for morphologies *n128*, *n172*, *n175* and *n184* respectively. **f, h, j.** Beeswarm plots (rectangles are medians) of percentage changes in *F*_max_ of the firing rate profiles obtained after virtual knockout of dendritic NaF (**f**), NMDARs (**h**) and KA channels (**j**) from each of the valid models across all morphologies. **g, i.** Beeswarm plots of percentage changes in FWHM of the firing rate profiles obtained after virtual knockout of dendritic NaF (**g**) and NMDARs (**i**) in valid models across all morphologies. **k.** Beeswarm plot of percentage changes in area under the curve (AUC) of the firing rate profiles obtained after virtual knockout of KA in valid models across all morphologies. The calculation of FWHM for models after virtual knockout of KA channels was not feasible because of spontaneous firing in KA knockout models.

### Computational details

All simulations were performed using the NEURON programming environment (Carnevale and Hines 2006), at 34**°** C with the resting membrane potential set at –65 mV. The simulation step size was 25 µs. Data analyses and graph plotting were performed using custom-written software in the IGOR Pro environment (Wavemetrics Inc.).

## RESULTS

Hippocampal CA1 neurons are complex and electrotonically non-compact structures with their dendritic terminals arranged in a laminar fashion (Mainen et al. 1996; Carnevale et al. 1997; Golding et al. 2005; Spruston et al. 1994). Both passive cable-filtering properties (Rall 1977; Stuart and Spruston 1998; Koch et al. 1983) and the non-homogenous distribution of active ion channels in the dendrites (Hoffman et al. 1997; Johnston et al. 1996; Johnston and Narayanan 2008; Magee 1998) contribute to differential attenuation of transient signals travelling both from dendrites to soma (Spruston et al. 1993; Golding et al. 2005) and vice-versa (Stuart and Sakmann 1994; Johnston et al. 1999). Although there is an overarching similarity in the gross morphology of CA1 pyramidal neurons, each neuron differs markedly in the structure of their dendritic trees. Given these differences and lines of evidence demonstrating the crucial roles of both dendritic geometry (Vetter et al. 2001; Krichmar et al. 2002; Narayanan and Chattarji 2010; Dhupia et al. 2014) and dendritic ion channel distributions (Narayanan and Johnston 2007, 2008, 2012; Spruston 2008; Magee 1999; Vaidya and Johnston 2013; Golding et al. 1999; Golding and Spruston 1998; Losonczy and Magee 2006; Hoffman et al. 1997; Spruston et al. 1995) in neuronal signal processing, it is imperative that the role of neuronal morphology on feature encoding be explored.

To do this, we looked at five different morphologically realistic reconstructions of hippocampal CA1 neurons and studied their place field encoding properties when identical inputs were fed to synapses afferent on each of them. We employed a conductance-based approach by distributing five different ion channel conductances in physiologically plausible gradients throughout the neuronal morphology and incorporated 100 excitatory synapses with randomly dispersed spatial locations. Each of these synapses was probabilistically activated by a Gaussian-modulated theta distribution function mimicking place field inputs, and across morphologies, each of these synapses received identical inputs. Specifically, whereas there was distinction (*i.e.* independent sampling of the same Gaussian-modulated theta distribution) in the inputs received by synapses within a neuronal morphology, synaptic input identity was maintained across morphologies (*e.g.*, synapse #1 of all morphologies received the same input, while inputs to synapse #1 were distinct from those to any other synapse). We then ran an exhaustive stochastic search, spanning thousands of parametric combinations, exploring several underlying intrinsic parameters in each morphology and analyzed the firing rate profiles of the models generated with these parametric/morphological combinations.

### Sharp place cell tuning was achieved across different morphologies, with disparate ion channel combinations and dispersed synaptic localization profiles

To quantify the structural differences in the five chosen CA1 pyramidal neuron morphologies, we ran Sholl’s analysis on them and found that they were distinct in terms of their size, dendritic length as well as their dendritic branching structure (Fig. 1). Specifically, there were heterogeneities in the overall branching structure and spread (Fig. 1a), the distribution of dendritic length (Fig. 1b) and branch points (Fig. 1c) across the apical and basal trees. In each of these five morphologies, we identified all possible distinct locations within the *stratum radiatum* as potential locations for synapse placement. Synapses were randomly placed at only 100 of these possible dendritic locations, of which there were 311–452 depending on the specific morphology under consideration (Fig. 1a). We confirmed that each of these potential synaptic locations had large pair-wise inter-synaptic distances in each of the five morphologies, ensuring that there was no spurious clustering as a consequence of randomized synaptic placements (Fig. 2a–b). We employed the base model parametric distributions of active and passive properties to set the synaptic permeability of AMPA receptors for synapses at each of these possible locations. These permeability values were location dependent (Fig. 2c) to set the somatic unitary EPSP values (Fig. 2d) at ∼0.2 mV (Basak and Narayanan 2018b; Narayanan and Chattarji 2010), consistent with electrophysiological observations that the somatic unitary EPSP values are location independent and are around ∼0.2 mV (Magee and Cook 2000; Smith et al. 2003; Andrasfalvy and Magee 2001). Together, whereas the choice of distinct reconstructions accounted for morphological heterogeneities in CA1 pyramidal neurons (Fig. 1), the randomized sampling of synaptic locations from all possible *stratum radiatum* locations incorporated heterogeneities in synaptic localization profiles (Fig. 2).

A third form of biological heterogeneity that is prevalent across CA1 pyramidal neurons pertains to ion channel conductance and their distribution profiles along the somatodendritic arbor (Hoffman et al. 1997; Magee 1998; Magee and Johnston 1995; Rathour and Narayanan 2012, 2014). To explore this parametric space (of 20 parameters) governing channel conductances and localization (Table 2), we employed an unbiased stochastic search paradigm that searched for models exhibiting sharp tuning to place field inputs. This search was performed on each of the five morphologies and for distinct synaptic localization profiles, thus ensuring that our search spanned structural and biophysical heterogeneities. Specifically, in employing the MPMOSS approach, we generated 8000–10000 models for each morphology, with each of the 20 parametric values randomly picked from independent uniform distributions spanning 0.5- to 2-fold of the base model values given in Table 2. The seed values of the randomization algorithm were kept constant during the stochastic-search run for each morphology, ensuring that the same 8000–10000 parametric combinations were tested across all five morphologies. The search space on ion channel conductances and their localization profiles was therefore *identical* across all the five morphologies that were tested.

We then activated the 100 randomly dispersed synapses with place-field inputs (Eq. 10) on each of these randomly generated models, recorded their voltage responses to these inputs (*e.g.*, Fig. 3A1) and calculated the FWHM and *F*_max_ for their firing rate responses (*e.g.*, Fig. 3A1). As the input activation distribution was common across all synapses, sharply tuned place cell responses were deemed to have low FWHM and high *F*_max_ (Basak and Narayanan 2018b), the cut-off for FWHM was set to be <2.8 s and that of *F*_max_ was set to be ≥ 40 Hz in all cases. We employed this relative approach, where we analyzed outputs with identical input distributions, for assessing tuning sharpness to ensure that our comparisons of the model responses remain focused on the biophysical and morphological heterogeneities. We obtained 2578/8000, 1065/10000, 4732/10000, 4205/10000 and 4573/10000 models for the morphologies *n123*, *n128*, *n172*, *n175* and *n184* respectively, that fulfilled these sharpness criteria. These results pointed to morphological characteristics of CA1 pyramidal neurons not being a constraining parameter in terms of encoding tuning sharpness.

### Expression of ion channel degeneracy in the concomitant emergence of sharp-tuning and physiologically relevant functional maps in the presence of morphological heterogeneities

Our analyses thus far had established the existence of several sharply tuned models in morphologically distinct neurons with randomly dispersed iso-feature synapses with randomly assigned channel condutances and their expression profiles. However, these models were not constrained by the electrophysiologically derived intrinsic properties of a CA1 pyramidal neuron. Would morphological heterogeneities be a constraining parameter if these sharply tuned place cells are additionally required to have physiologically relevant intrinsic properties? To answer this question, we subjected the sharply tuned models in all the five morphologies to a further step of validation involving intrinsic functional maps (Narayanan and Johnston 2012), where the measurements of four intrinsic functional maps were validated against their respective electrophysiological bounds mentioned in Table 3. This second step of validation yielded 152, 42, 136, 412 and 90 *valid* models for the morphologies *n123*, *n128*, *n172*, *n175* and *n184* respectively, which concomitantly had sharply tuned place cell response and electrophysiologically validated intrinsic functional characteristics. Of these valid models, we chose five examples from each morphology that exhibited similar tuning properties (FWHM and *F*_max_; *e.g.*, Fig. 3A1 for morphology n123) *and* similar intrinsic functional maps (*e.g.*, Fig. 3A2–A3). We noted similarity in tuning sharpness *and* in intrinsic measurements across the distinct morphologies that we had employed (Fig. 3). These results suggest that morphological heterogeneities constitute a sloppy, and not a stringent parameter, in the concomitant emergence of place cell encoding and robust intrinsic functional maps.

In CA1 pyramidal neurons, ion channel degeneracy has been previously demonstrated with regard to the emergence of intrinsic neuronal properties (Rathour and Narayanan 2012; Srikanth and Narayanan 2015; Migliore et al. 2018), intrinsic functional maps (Rathour and Narayanan 2014; Rathour et al. 2016), short- (Mukunda and Narayanan 2017) and long-term (Anirudhan and Narayanan 2015) synaptic plasticity profiles, and the concomitant emergence of place cell encoding and intrinsic functional maps (Basak and Narayanan 2018b). However, the role of diverse neuronal morphology has not been closely examined in the context of place cell encoding within the ion channel degeneracy framework. Do disparate combinations of ion channels lead to similar sharpness in place field response while exhibiting signature electrophysiological characteristics in neuronal models built with distinct morphological substrates? To answer this, we first looked at the distribution of parameters underlying the five similar valid models derived from each of the five morphologies (5 morphologies × 5 examples each). As an example, Fig. 3A4 depicts the normalized parametric distributions of the five example valid cells (from Figs. 3A1–A3) with similar sharp tuning as well as analogous intrinsic properties corresponding to the morphology *n123*. We noted that the parameters underlying these similar models were not similar and did not exhibit clustering around a specific value, but spanned a broad range of their respective domains (Fig. 3A4). This observation was true not just for mophology *n123*, but extended to other morphologies as well (Fig. 3b–e).

To ensure that this finding was not limited to only the five example models, we explored tuning sharpness (Fig. 4), the electrophysiological measurements (Fig. 4) and the underlying parametric distributions (Fig. 5) of all the valid cells across each of the five morphologies. We confirmed that the tuning sharpness of models from each of the five morphologies (Fig. 4a–b) and the intrinsic measurements that represented the four functional maps (Fig. 4c–f) were within their respective bounds and were similar across all five morphologies. Next, we assessed the individual distributions of all 20 underlying MPMOSS parameters in all valid models for each morphology (Fig. 5a). Consistent with our observations with the five selected models (Fig. 3), we found the individual parametric histograms to span a broad range of their respective assigned ranges (see Table 2) for all morphologies (Fig. 5a).

The analyses thus far showed that there was no need for individual channels to maintain specific expression or localization profiles, and that several combinations of dispersed synaptic localization and disparate channel localization profiles are capable of eliciting sharp tuning despite morphological differences. Were there pairwise compensations in these parameters, whereby expressions of different channels were correlated in a pairwise manner? To address this, we plotted the pairwise correlation matrix (Pearson’s correlation coefficient) spanning all the 20 parameters of valid models from each of the five morphologies (Fig. 5b). These matrices revealed weak pairwise correlations (statistics of correlation coefficients for the different morphologies: *n123*: max=0.42; min=–0.6; mean ± SEM= 0.02 ± 0.006; *N*_valid_ = 152. *n128*: max= 0.54; min= –0.7; mean ± SEM= –0.006 ± 0.01; *N*_valid_ = 42. *n172*: max= 0.48; min= –0.55; mean ± SEM= 0.007 ± 0.006; *N*_valid_ = 136. *n175*: max= 0.53; min= –0.46; mean ± SEM= 0.005 ± 0.005; *N*_valid_ = 412. *n184*: max= 0.49; min= –0.55; mean ± SEM= 0.013 ± 0.007; *N*_valid_ = 90) between the valid-model parameters across all morphologies (Fig. 5b).

Across morphologies, we employed the same parametric search space governing channel expression and localization, and assessed the impact of morphological differences employing *identical* parametric values across all morphologies. This experimental design allowed us to ask if *identical* parametric values resulted in valid models with different morphologies, and thereby delineate the role of morphological sloppiness from that of ion channel degeneracy. Taking advantage of this experimental design, we listed unique parametric combinations that resulted in valid models only in any one of the five morphologies, and those parametric combinations that resulted in valid models in more than one morphology. Of the total 832 valid parametric combinations spanning all morphologies, we found a majority of models (727) to yield valid models for only one of the five morphologies. However, importantly, we found 87 valid-model parametric combinations common to two morphologies, 14 combinations common to three morphologies and 4 combinations common to four morphologies (Fig. 5c). We did not find any valid-cell parametric combination that was common to all five morphologies. These results provide strong lines of evidence that morphological heterogeneities do not impose strong constraints in the expression of degeneracy in sharp place-field encoding and the concomitant emergence of signature intrinsic functional maps in hippocampal pyramidal neurons.

### Spatially dispersed synapses yielded dendritic spikes in all morphologies

Previous studies have demonstrated the importance of dendritic spikes (dSpikes) in sharp place cell responses (Sheffield and Dombeck 2015; Basak and Narayanan 2018b). Do dendritic spikes play a differential role in sharp place-field tuning based on the heterogeneities in neuronal morphology? To answer this, we assessed the dynamics of dSpike generation in the valid cells across all morphologies by simultaneously recording voltage traces at various locations along the somato-apical trunk (Fig. 6a), in response to place cell input through randomly dispersed synaptic locations. We analyzed these traces to quantify the proportion of dSpikes generated at each of the recording locations in each cell (Basak and Narayanan 2018b). Specifically, we compared the timing of the depolarization at these locations corresponding to each somatic AP. If the dendritic peak voltage preceded the somatic AP, we considered it to be an indication of the generation of a dSpike, whereas if the dendritic peak followed the somatic AP, we considered it to be a backpropagating action potential (bAP). We then collated all the timing data at each location from each valid cell in each morphology and plotted location-dependent time-difference histograms (Fig. 6b). Negative time differences in these histograms indicated the generation of dSpikes whereas positive time differences were representative of bAPs. We noted that a substantial fraction of somatic spikes were preceded by dSpikes across all morphologies (Fig. 6b). Although *n184* shows comparatively lesser amount of dSpikes quantifiable from voltage recordings along the dendritic trunk, we confirmed that dendritic spikes were highly prevalent in an apical oblique of morphology *n184* (Fig. 6b, inset on far right panel; dSpike fraction=0.57). Together, although there was heterogeneity in the relative distribution of dendritic spikes at different dendritic locations, these observations clearly demonstrated that the generation of dendritic spikes with spatially dispersed synapses was prevalent across all morphologies.

### Bidirectional modulation of dendritic spike prevalence revealed the critical role of dendritic spikes in tuning sharpness of place field responses across all morphologies

How does the presence of these dendritic spikes affect sharp tuning of place-field responses? How would sharpness of tuning change in these models if dendritic spikes generation were suppressed or enhanced by changes in channel/receptor properties? Do different intrinsic parameters have differential effect on dSpike generation in neurons owing to their morphological differences? To introduce bidirectional changes in dendritic spike prevalence, we employed the virtual knockout model (VKM) approach (Rathour and Narayanan 2014; Basak and Narayanan 2018b; Mukunda and Narayanan 2017; Mittal and Narayanan 2018) on each of the valid models across all five morphologies. The VKM approach offers an ideal route to understand the implications of individual channels and receptors in a heterogeneous population of models exhibiting degeneracy. In this approach, to assess the effect of a particular conductance or receptor, we set that channel conductance or receptor permeability value to zero, recorded the model voltage and firing responses to identical synaptic inputs and compared them to their corresponding baseline model (where the channel/receptor was intact). Motivated by electrophysiological and imaging data (Johnston et al. 2000; Losonczy and Magee 2006; Golding et al. 1999; Sheffield and Dombeck 2015; Hoffman et al. 1997; Gasparini et al. 2004) and prior computational models of place cells (Basak and Narayanan 2018b), we employed virtual knockout of dendritic NaF channels (Fig. 6c) or NMDA receptors (Fig. 6d) for reducing the prevalence of dendritic spikes and virtual knockout of KA channels (Fig. 6e) to enhance their prevalence.

As expected, knocking out either dendritic NaF (Fig. 6c, Fig. 7a–e) and NMDARs (Fig. 6d, Fig. 7a–e) resulted in a substantial reduction in dSpike fraction across all morphologies. The effect was more prominent for the dNaF knockouts, as this conductance constitutes the predominant regenerative depolarizing force required to sustain dSpikes (Golding and Spruston 1998; Losonczy and Magee 2006). As a consequence to the loss of dSpikes in dNaF or NMDAR virtual knockouts, models across all morphologies showed a substantial loss in tuning sharpness (Fig. 7f–i). Virtual knockout of KA channels drastically increased the prevalence of dSpikes across all morphologies (Fig. 6e, Fig. 7a–e). Owing to an expected (Losonczy and Magee 2006; Frick et al. 2003; Hoffman et al. 1997; Johnston et al. 2003; Johnston et al. 2000; Chen et al. 2006; Kim et al. 2005; Rathour et al. 2016) increase in intrinsic excitability after removal of the KA conductance, KA VKMs exhibited an increased firing rate (Fig. 7j). As a consequence, some of the models spontaneously fired throughout the recording period (even outside the place field input period), and it was difficult to quantify the FWHM of their firing rate profiles. Therefore, we calculated the percentage change in the area under the curve (AUC) of the firing rate profile in the KA^−^ case across all morphologies (Fig. 7k). Both *F*_max_ and AUC increased in the KA VKMs for all morphologies, indicating a drastic loss of sharpness in tuning to place-field inputs, where cells fired even in the absence of place field inputs. Together, whereas a reduction in dendritic spike prevalence resulted in loss of tuning sharpness through loss of overall depolarizing drive, an enhancement in dendritic spike prevalence accompanied by a non-specific enhancement in excitability also resulted in loss of tuning sharpness. These results clearly show that sharpness in feature tuning is maintained by an intricate balance between mechanisms that promote and those that prevent dendritic spike generation. These results also demonstrate that the mechanism behind the dynamics of dSpike generation stay consistent across different morphologies.

## DISCUSSION

The main conclusion from this study is that sharp place-field tuning is achieved in hippocampal CA1 pyramidal neurons with randomly dispersed iso-feature synaptic inputs irrespective of geometrical heterogeneities in the underlying morphologies. These analyses establish heterogeneities in neuronal morphology to be a sloppy, and not a stringent parameter from the perspective of the concomitant emergence of sharp feature encoding and signature electrophysiological characteristics. We demonstrated this using an unbiased stochastic search approach that spanned several biophysical properties and distinct patterns of random dispersion of iso-feature synapses across several morphologies. The sloppy nature of morphological characteristics from an electrophysiological standpoint, especially in an electrotonically non-compact neurons was actively mediated by the presence of somato-dendritic channel expression. Importantly, we show that the concomitant emergence of several intrinsic functional maps along with sharp feature tuning are achievable with not one specific expression profile of ion channels and receptors, but with disparate combinations of channel expression and randomly dispersed synaptic localization profiles. Thus dispersed synaptic localization and ion channel degeneracy together compensate for neuron-to-neuron variability in neuronal geometry in maintaining the electrotonic structure, in the emergence of intrinsic functional maps and sharp feature tuning. Employing these morphologically and biophysically disparate, yet functionally similar models as substrates, we explored the impact of bidirectional modulation in dendritic spike prevalence by virtual knockout of specific channels and receptors. We found that knock out of dendritic sodium channels (dNaF) or NMDA receptors reduced the prevalence of dendritic spikes, whereas knockout of transient potassium (KA) channels enhanced dendritic spike prevalence across all tested morphologies. However, irrespective of which of these three channels/receptor were knocked out, tuning sharpness reduced owing either to an overall reduction in the depolarizing drive (NMDAR/dNaF knockout) or due to a non-specific enhancement in intrinsic excitability (KA knockout). These results further emphasize the need for synergy and balance between regenerative and restorative conductances in maintaining dendritic spike prevalence as a tightly regulated physiological variable and in ensuring tuning sharpness and excitability homeostasis. Neuronal tuning sharpness can, therefore, be altered through neuromodulation, through activity-dependent plasticity or through channelopathies that alter dendritic spike prevalence.

### Strategy for concomitant sharp feature encoding and robust intrinsic functional maps in neurons exhibiting morphological heterogeneity

The impact of neuronal morphology has been explored from several perspectives, and could be placed under at least three distinct classes. The first class of studies have explored the physiological implications of qualitative morphological differences across *different* neuronal subtypes (Mainen and Sejnowski 1996; Cannon et al. 2010; Vetter et al. 2001). A second class of studies have assessed pathology-induced or behaviorally relevant dendritic remodeling (atrophy or hypertrophy) in neuronal structures (van Elburg and van Ooyen 2010; van Ooyen et al. 2002; Narayanan and Chattarji 2010; Narayanan et al. 2005; Dhupia et al. 2014). A third class of studies explored the implications for intrinsic morphological heterogeneities *within* a neuronal subtype (Krichmar et al. 2002; Ferrante et al. 2013; Beining et al. 2017; Weaver and Wearne 2008; Otopalik et al. 2017b; Otopalik et al. 2019; Otopalik et al. 2017a; Schaefer et al. 2003). These studies have assessed several functional outcomes, with the focus on overall excitability characteristics (input resistance, firing rate, firing patterns, impedance characteristics including resonance) and propagation characteristics (somato-dendritic coupling, dendritic spike initiation and propagation, backpropagation of action potentials). Although the specifics of the overall findings were dependent on the function that was being assessed, a general conclusion across these studies is that neuronal morphological properties play a critical role in modulating function, when *all other components are set to identical values.* For instance, when somato-dendritic channel distributions are maintained to be identical, morphological differences contribute to differences in firing rate, firing pattern, backpropagation of action potentials and the emergence of several functional maps along the somato-dendritic arbor (Mainen and Sejnowski 1996; Vetter et al. 2001; Narayanan and Chattarji 2010; Dhupia et al. 2014).

However, recognizing neuronal morphology to be one of the several variables that regulate function, studies have also shown that functional changes introduced by morphological remodeling could be compensated by appropriate changes in other components. For instance, although dendritic atrophy was shown to increase firing rate and reduce burst propensity in CA3 pyramidal neurons, these changes in *atrophied neurons* could be reversed by altering the conductance densities of one of several ion channels expressed in CA3 pyramidal neurons (Narayanan and Chattarji 2010). Within this overall thesis of disparate biophysical components compensating for morphological differences, our study directly demonstrates that conjunctive changes in *several* combinations of channels could yield similar functional outcomes *despite* the presence of heterogeneous morphological characteristics. Our study quantitatively demonstrates this possibility for the concomitant emergence of sharp tuning characteristics along with several intrinsic functional maps.

This demonstration was contingent on synaptic democracy, on the ability of randomly dispersed synapses to yield sharp tuning through generation of dendritic spikes, and on ion channel degeneracy. Synaptic democracy (Magee and Cook 2000; Andrasfalvy and Magee 2001; Smith et al. 2003; Hausser 2001) ensured that the electrical impact of somatodendritic excitatory synapses on the soma remained the same irrespective of their location in the dendritic tree, thereby laying the substrate for sharp tuning with randomly dispersed iso-feature synapses in morphologically heterogeneous structures (Figs. 2–4). This necessity implied that the synaptically evoked voltages at the synaptic location were large, consistent with experimental observations (Harnett et al. 2012; Jayant et al. 2017; Kwon et al. 2017; Popovic et al. 2015), and provided the depolarizing drive essential for the generation of dendritic spikes (Basak and Narayanan 2018b). The presence of dendritic ion channels and associated degeneracy enabled the emergence of several intrinsic functional maps (Basak and Narayanan 2018b; Rathour and Narayanan 2014), and the generation of dendritic spikes ensuring temporal precision of somato-dendritic information transfer (Basak and Narayanan 2018b) and sharpness of feature tuning across in morphologically heterogeneous structures (Figs. 6–7). Together, our study proposes the recruitment of synaptic democracy and ion channel degeneracy towards eliciting dendritic spikes through dispersed synaptic localization as an effective strategy for achieving concomitant sharp feature encoding and robust intrinsic functional maps in neurons exhibiting morphological heterogeneity. This strategy provides neurons with several degrees of freedom in terms of the specific localization of synapses, the morphological micro-structure and the expression profiles of different ion channels. Importantly, as a consequence of the randomly dispersed nature of iso-feature synapses and morphological heterogeneity being compensated by other biophysical characteristics, this strategy precludes the need for *precise* wiring to achieve precision in the functions considered in this study.

### Morphological sloppiness in electrotonically non-compact neurons

A set of previous studies explored the role of morphological heterogeneity on functional outcomes in STG neurons (Otopalik et al. 2017b; Otopalik et al. 2019; Otopalik et al. 2017a), which are required to generate slow temporal network dynamics. These studies demonstrated that STG neurons were able to achieve equivalent functional outcomes in several morphologically distinct neurons as a consequence of their *electrotonically compact* structure and slow voltage dynamics, which are relatively resilient to attenuation (Rall 1977). In this study, we have demonstrated sloppiness of morphological heterogeneity in an *electrotonically non-compact* neuronal subtype, that is required to sustain the propagation of fast events including dendritic spikes and backpropagating action potentials (Spruston 2008; Spruston et al. 1995; Spruston et al. 2007; Gasparini et al. 2004; Golding et al. 1999; Golding et al. 2005; Golding and Spruston 1998). One of the specific requirements that mediates the sloppy nature of morphological heterogeneities in electrotonically non-compact neuronal structures is that the electrotonic characteristics of the diverse morphological structures are similar. In our analyses, this specific requirement for sloppiness of morphology in electrotonically non-compact neurons is ensured by the ability of these neuronal substrates to exhibit several intraneuronal functional maps (especially backpropagating action potential attenuation, and attenuation of EPSP propagation in the maintenance of synaptic democracy) through ion channel degeneracy. This ensures that the somatic and dendritic compartments are electrotonically distant, providing the substrate for generation of dendritic spikes that may not propagate to generate a somatic action potential (Gasparini et al. 2004; Golding et al. 1999; Golding et al. 2005; Golding and Spruston 1998; Losonczy and Magee 2006; Moore et al. 2017; Basak and Narayanan 2018b).

A central requirement here is the ability to maintain several intraneuronal functional maps, which in turn ensures that the active electrotonic structure of the neuronal morphology is maintained. In our analyses, this requirement is fulfilled by ion channel degeneracy, which in conjunction with the synaptic democracy and randomly dispersed synapses ensured that the morphological microstructure was not a strongly constraining stringent parameter in achieving the specific functional goals assessed here. Together, in electrotonically compact neurons, animal-to-animal variability in neuronal geometry was shown to be compensated for by compact electrotonic structure (Otopalik et al. 2017b; Otopalik et al. 2019; Otopalik et al. 2017a). In electrotonically non-compact neurons, for the specific function of eliciting tuning sharpness and maintaining intraneuronal functional maps, dispersed synaptic localization and ion channel degeneracy together could compensate for neuron-to-neuron variability in geometry. In extending our conclusions to other electrotonically non-compact neuronal subtypes, it is important to account for the extent of morphological heterogeneity, the ability of ion channel degeneracy to maintain electrotonic structure and intrinsic functional maps, the specific function that is under consideration and the other components that interact with morphology in eliciting the specific function. Importantly, future studies could explore the systematic morphological, transcriptional and electrophysiological heterogeneities across the dorso-ventral, superficial-deep and proximal-distal axes of the hippocampus (Malik et al. 2016; Dougherty et al. 2012; Igarashi et al. 2014; Cembrowski and Spruston 2019; Soltesz and Losonczy 2018; Cembrowski et al. 2016; Strange et al. 2014). From the functional standpoint of this study, such future studies could ask if these morphological heterogeneities significantly contribute to changes in tuning sharpness for place fields observed there and somatodendritic functional maps in those neurons, or if these heterogeneities are non-constraining variables compensated for by interactions with other components that are expressed there.

### The impact of neuronal morphology on neuronal physiology is function-specific and is critically reliant on interactions with several other components

The specific function under consideration and the role of the interactional space encompassing other components cannot be overemphasized in the assessment of the impact of neuronal morphology on physiology. Our conclusion on the sloppy nature of morphological heterogeneities in arriving at sharp feature tuning and robust intrinsic electrophysiology was aided by synaptic democracy, by the ability of randomly dispersed synapses to yield sharp tuning, and by ion channel degeneracy. The absence of these *other components*, and their *interactions* with the neuronal morphology under consideration, would have rendered our conclusion on sloppiness infeasible. Additionally, while our assessment on sloppiness holds for achieving the *specific functions* of sharp place-field tuning and signature electrophysiological characteristics, these analyses would not necessarily hold if another functional metric were to be chosen.

An example for such a function where morphological characteristics and associated heterogeneities would have be assessed at a different spatial resolution and might play a stronger role in altering function is the generation and propagation of signaling microdomains. The generation and propagation of signaling microdomains along a biochemical network is at the center of several location-specific plasticity paradigms, and are known to be dependent on neuronal morphological characteristics such as neuronal diameter (Ashhad and Narayanan 2013; Basak and Narayanan 2018a; Berridge 2006; Neves and Iyengar 2009; Neves et al. 2008). Importantly, the length constants associated with the propagation of these signaling microdomains are orders of magnitude less than the propagation of voltage signals (Zador and Koch 1994; Augustine et al. 2003; Rizzuto and Pozzan 2006; Basak and Narayanan 2018a). For instance, the length constant associated with calcium microdomains are less than a micron, whereas the electrotonic length constant for neuronal processes is on order of 100s of microns (Zador and Koch 1994). Thus, although spatially dispersed synapses endowed with synaptic democracy could interact across large distances, owing to the large electrotonic distances, to yield sharp feature tuning, their calcium microdomains would not interact to yield a neuron-wide somatic spiking response. In this scenario, the local dendritic diameter, the dendritic branching patterns, the motif of the biochemical network and the distribution profiles of the several molecular species would play critical roles in regulating the generation and propagation of signaling microdomains (Basak and Narayanan 2018a; Neves and Iyengar 2002, 2009). Importantly, heterogeneities in morphological characteristics could play a much larger and severely constraining role in such a scenario, although they were not important from an electrotonic perspective. An ideal example for this distinction is on electrical *vs.* biochemical compartmentalization (Koch and Zador 1993) in dendritic spines (*e.g.*, confinement of calcium signals within a spine owing to the neck acting as a diffusion barrier).

It is therefore important to ensure that generalizations are avoided in describing the impact of neuronal morphology on physiology, and the observations be confined to specific functions that are being assessed in a study while also emphasizing the importance of other components in the emergence of these functions. While morphological heterogeneities might not be a strong constraining parameter in achieving one physiological goal, these heterogeneities might be the defining parameter for another physiological attribute of the neuron, in a manner that is critically reliant on other components as well.

Together, the broad framework where ion channel degeneracy and randomly dispersed iso-feature synapses coexist, morphological heterogeneities are flexible and sloppy parameters that don’t impose stringent constraints on feature encoding or intrinsic electrophysiological characteristics. Ion channel degeneracy, the sloppy nature of morphological characteristics and the dispersed nature of iso-feature synapses together provide neurons with significant flexibility in terms of network wiring and channel/receptor localization profiles that would elicit sharp feature encoding along with robust excitability homeostasis.

## AUTHOR CONTRIBUTIONS

R.B. and R. N. designed experiments; R.B. performed experiments and carried out data analysis; R.B. and R. N. co-wrote the paper.

## ACKNOWLEDGMENTS

The authors thank members of the cellular neurophysiology laboratory for helpful discussions and for comments on a draft of this manuscript.

